# Statistical modeling and analysis of multiplexed imaging data

**DOI:** 10.1101/2023.03.17.533200

**Authors:** Pierre Bost, Ruben Casanova, Uria Mor, Bernd Bodenmiller

## Abstract

The rapid development of multiplexed imaging technologies has enabled the spatial cartography of various healthy and tumor tissues. However, the lack of adequate statistical models has hampered the use of multiplexed imaging to efficiently compare tissue composition across sample groups, for instance between healthy and tumor tissue samples. Here, we developed two statistical models that accurately describe the distribution of cell counts observed in a given field of view in an imaging experiment. The parameters of these distributions are directly linked to the field of view size and also to properties of the studied cell type such as cellular density and spatial aggregation. Using these models, we identified statistical tests that have improved statistical power for differential abundance testing of tissue composition compared to the commonly used rank-based test. Our analysis revealed that spatial aggregation is the main determinant of statistical power and that to have sufficient power to detect differences in cell counts when cells are highly aggregated may require sampling of hundreds of fields of view. To overcome this challenge, we provide a new stratified sampling strategy that might significantly reduce the number of required samples.

## Introduction

Efficient and robust data analyses rely on a correct understanding of the properties of the studied data. Identification of the most suitable probabilistic model for a given data type can be cumbersome, but once identified, probabilistic modeling dramatically improves the quality of data analysis [Lisitsin et al., Brill et al.]. A prime example comes from the field of single-cell RNA sequencing data analysis where different experimental protocols generate data with different statistical properties [Svensson], thus requiring that different statistical models be used to perform routine analysis such as differential gene expression or differential abundance analyses.

We reasoned that this would also be true for multiplexed imaging datasets, which involve measurements of dozens to thousands of features with a cellular to sub-cellular resolution [Moffitt et al.]. Multiple techniques have been developed to measure abundances and localizations of transcripts [Shah et al., Eng et al.], proteins [Giesen et al., Goltsev et al., Moffit et al., Hickey et al.], and epigenome marks [Lu et al.]. Multiplexed imaging has been used to dissect the biology of diverse healthy tissues [Shah et al.] and the pathogenesis of various diseases [Rendeiro et al., Jackson et al.], but no dedicated statistical models have been developed to analyze multiplexed imaging data, and most published analyses rely on non-parametric tests, potentially reducing sensitivity to detect changes in tissue composition and hampering the potential of biological discovery [Blair et al.].

Considerable work has previously been done to develop methods for analysis of spatial patterns in the fields of ecology [Wiegand et al.], forestry [Penttinen et al.], and mineralogy [Lisitsin et al.], resulting in the development of spatial point pattern theory (SPPT) and efficient analysis algorithms [Illian et al]. SPPT and its associated tools have rarely been used to analyze cellular biology spatial data, however [Edsgärd et al.], and most work in the biological realm has focused on the analysis of low-dimensional imaging data generated using immunofluorescence or immunohistochemistry [Rockhill et al., Diggle et al]. These studies did not model cell counts within biological tissue, which must be done to correctly estimate the spatial density of a given cell type and to quantify differences in cell composition between samples.

Here we introduce new statistical models inspired by the SPPT to model the distribution of cell counts across fields of view (FoVs) in multiplexed imaging data. Our models were validated on datasets generated by imaging mass cytometry (IMC) [Giesen et al.]. We tested the ability of statistical tests based on those models to detect slight differences in cellular density and observed a strong correlation between cell spatial pattern and statistical power. Tests based on our models outperformed the non-parametric rank test for most studied cell types, and a large number of samples were needed for accurate quantification of rare and highly spatially aggregated cell phenotypes. We addressed this challenge by introducing a new sampling strategy inspired by stratified sampling [Cochran]. This approach drastically decreased the number of samples needed to perform differential abundance analysis between sample groups and is broadly applicable to any type of multiplexed imaging experiment.

## Main text

### Identification of adapted statistical models for multiplex imaging data

In order to identify a statistical model able to accurately model the distribution of cell counts across FoVs (also known as quadrat counts), we analyzed our previously published dataset of a healthy human lymph node imaged using IMC [Bost et al.]. We computationally sampled 200 square FoVs of 400 µm width and counted the numbers of cells present for each of the 17 different cell phenotype clusters we previously identified [Bost et al.]. We fitted three discrete probability distributions to the observed cell count data for each individual cell phenotype cluster (Figure 1A):

**Figure 1:**
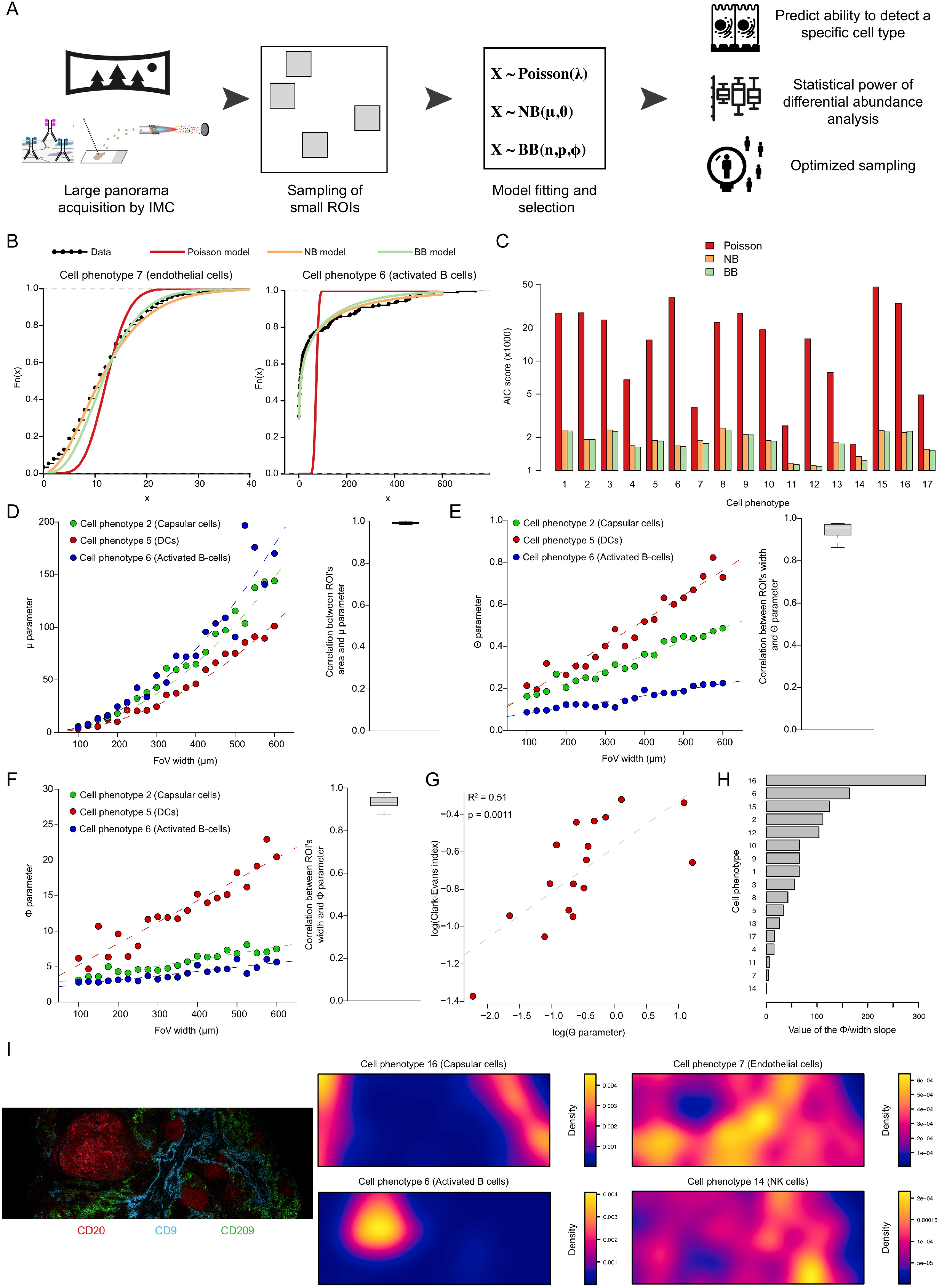
Modeling multiplexed imaging cell count data using beta and negative binomial distribution. **(A)** Analytical workflow. **(B)** Empirical cumulative distribution function of quadrat counts (black line) for cell phenotypes 7 (left) and 6 (right) identified in healthy human lymph node by Bost et al. compared to the theoretical cumulative distribution functions of the three fitted theoretical distributions (red: Poisson, orange: negative binomial (NB), green: beta binomial (BB)). **(C)** AIC scores of the three fitted models for each cell phenotype identified previously. **(D)** Left: Relationship between FoV width and negative binomial **µ** for three representative cell phenotypes. The dashed lines correspond to quadratic fits. Right: Distribution of R^2^ values for the quadratic relationship between FoV width and **µ** across all cell phenotypes. The thick line corresponds to the median, and the bottom and upper limits of the box correspond to the first and third quartiles, respectively. The lower and upper whiskers correspond to the lowest and highest values, respectively, within the range of the first and third quartiles ±1.5 times the interquartile range (IQR). **(E)** Left: Relationship between FoV width and negative binomial parameter **Θ** for three cell phenotypes. Dashed lines correspond to the linear model between FoV width and **Θ**. Right: Distribution of R^2^ values for the linear relationship between FoV width and the **Θ** across all cell phenotypes. **(F)** Left: Relationship between FoV width and beta binomial parameter **ϕ** for three cell phenotypes. Dashed lines correspond to the linear model between FoV width and **ϕ**. Right: Distribution of R^2^ values for the linear relationship between FoV width and **ϕ** parameter across all cell phenotypes. **(G)** Association between the negative binomial parameter **Θ** and the Clark-Evans index. The dashed line corresponds to the linear regression linking the two variables. **(H)** Ranked values of the slope between the width and ϕ for each cell phenotype. **(I)** Left: Tissue structure of the lymph node sample imaged by IMC. Right: Spatial densities of four cell phenotypes across the tissue.

- The Poisson distribution is parametrized by a single parameter **λ** corresponding to its mean and variance, which are assumed to be equal thereby limiting its flexibility. This distribution is commonly used to model quadrat counts of spatially homogenous point patterns [Illian et al.].
- The negative binomial distribution is an extension of the Poisson distribution where the parameter **λ** is randomly sampled from a gamma distribution defined by two parameters, the mean **µ** and the over-dispersion parameter **Θ**, the latter parameter dealing with high variability not accounted for in a Poisson model. This distribution has been used to model quadrat counts of aggregated point patterns despite the absence of theoretical point patterns exhibiting such a quadrat count distribution [Diggle].
- The beta binomial distribution is an extension of the binomial distribution where the probability of success is randomly sampled for a beta distribution. The distribution is defined by the total number of trials **n**, the mean probability of success **p**, and an over-dispersion parameter **ϕ**. This distribution also results in over-dispersion, but it differs from the negative binomial distribution in that its values are bounded by the **n** parameter, a key feature that results in realistic modeling of quadrat counts. The distribution assumes that only a bounded number of cells can be present in a FoV.

Based on Akaike Information Criterion (AIC) scores, which provide an estimate of a model quality considering the model likelihood and number of parameters, both the negative and the beta binomial distributions systematically outperformed (i.e., had lower AIC scores than) the Poisson distribution for every individual cell phenotype in the IMC dataset (Figure 1B and C). This is because only spatially homogeneous point patterns can generate quadrat count data that follow a Poisson distribution [Illian et al. 2008], and distributions of most cell types in lymph nodes are not to be homogenous. While the beta-binomial distribution usually fitted better the sampled data than the negative binomial distribution, the difference between the two models was small (for 15 out of 17 cell phenotypes, Figure 1C). We then investigated the impact of the FoV size on the parameters of the fitted negative and beta binomial distributions by sampling FoVs of widths ranging from 100 µm to 600 µm. We observed that the mean **µ** parameter of the negative binomial distribution that corresponds to an individual cell phenotype is proportional to the FoV area and that each cell phenotype has a different slope (Figure 1D). In addition, the **Θ** over-dispersion parameter scaled linearly with the FoV width, and each cell phenotype analyzed had a different slope and intercept (representative cell phenotype clusters shown in Figure 1E). Similarly, the beta binomial **ϕ** over-dispersion parameter scaled linearly with FoV width, and each cell phenotype had a unique slope and intercept (Figure 1F). As both **Θ** and **ϕ** parameters directly control over-dispersion, larger FoVs will result in less noisy quadrat counts and more robust cell density estimations.

We then investigated whether over-dispersion parameters could be linked to distinct spatial patterns. We observed a weak but significantly positive correlation between the transformed over-dispersion parameter **Θ** of each cell phenotype and its respective Clark-Evans index (Figure 1G), a crude measure of spatial aggregation based on nearest-neighbor distance (Illian et al.). This positive correlation indicates that a highly spatially aggregated point pattern (i.e., low Clark-Evans index) tends to yield highly over-dispersed count data (i.e., low over-dispersion parameter). Lastly, we identified cell phenotypes with the highest and lowest ratios of the over-dispersion parameter to FoV width and asked whether these phenotypes differ in their spatial distributions. We observed that those cell phenotypes with steepest slopes, capsular and activated B cells, were localized in specific regions of the lymph node, whereas those with flatter slopes, natural killer (NK) and endothelial cells, were homogeneously spread across the tissue (Figure 1H and I). This validated our interpretation of the over-dispersion parameters as a measure of the spatial aggregation of the data.

To determine whether these findings are generalizable, we analyzed IMC datasets from two human melanoma samples, each with a different immune infiltration pattern (Figure S1A and B). The three aforementioned models were fit to the data, and conclusions were similar to those reached by analysis of the lymph node dataset. The Poisson model gave higher AIC scores than the two binomial distribution models for both melanoma samples (Figure S1C and D). The AIC scores for the binomial distribution models were similar for most cell phenotypes (Figure S1C and D). In addition, a linear relationship between the FoV width and the fitted **Θ** parameter for the negative binomial model was observed in data for both samples (Figure S1E and F) as was the linear relationship between the FoV width and the beta binomial distribution **ϕ** parameter (Figure S1G and H). These analyses demonstrated that it is possible to model multiplexed imaging quadrat count data using the over-dispersed negative and beta binomial distributions and that parameters of these models are directly linked to the studied cell phenotype spatial pattern and FoV size.

### Negative and beta binomial distribution models recapitulate basic properties of random sampling

We recently developed an empirical model that describes the probability of detecting a given number of cell phenotypes when sampling a set number of FoVs within a biological tissue [Bost et al]. The relationship between the number of imaged FoVs, r, and the number of recovered cell phenotypes, N(r), was defined by the following equation:

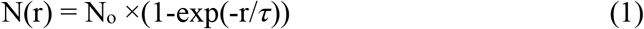

where N_o_ is the total number of cell phenotypes present in the tissue, and *τ* indicates how many regions must be imaged to recover most known cell phenotypes (i.e., 2*τ* FoVs must be imaged to detect more than 85% of the cell phenotypes). This model assumed that the cell phenotype clusters are spatially homogenous within a tissue given the quadrat count follows a Poisson distribution. Furthermore, *τ* was sensitive to a cell recovery threshold **T**, which describes the minimal number of observed cells that must be recovered to determine if a cell phenotype has been detected [Bost et al.]. The models presented here overcome these limitations. We fitted negative and beta distribution models of each individual cell phenotype with various spatial aggregation from the lymph node sample to compute the probability of detecting each phenotype within a given number of FoVs of 400 µm width. Both models perfectly predicted the empirical estimations of the *τ* parameter (Figure 2A, R^2^>0.99 for both models), showing that phenotypes of various spatial aggregation can be modeled. We then used the negative binomial distribution model to investigate the detection of individual cell phenotypes using the cell recovery threshold parameter **T**. The relationship between **T** and *τ* was asymptotically linear across all cell phenotypes (Figure 2D). Each cell phenotype had a different linear slope, demonstrating that the chosen value of **T** impacts the detection of each cell phenotype differently. As the probability of detecting individual (and especially rare) cell phenotypes can be precisely computed using either binomial distribution model with high precision, the researcher can select the optimal experimental design necessary to observe and characterize a particular rare cell phenotype (Figure 2E). In sum, the binomial distribution models perfectly recapitulate empirical sampling analysis and can be used to improve the design of multiplexed imaging experiments to detect individual cell phenotypes within a tissue.

**Figure 2:**
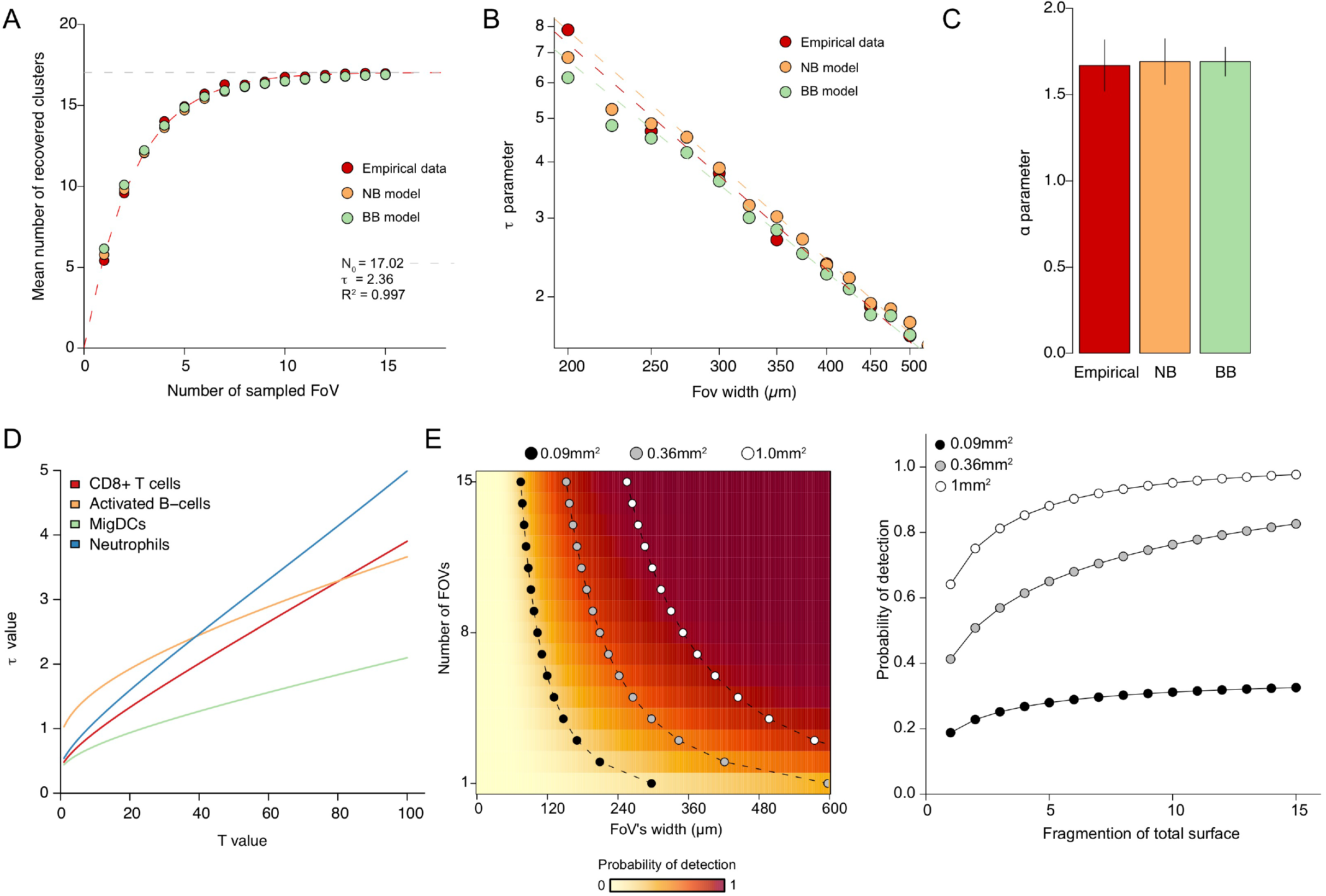
Prediction of cell phenotype recovery probability using the binomial models. **(A)** Theoretical and empirical mean numbers of cell phenotypes recovered from a given number of FoVs. The red dashed line corresponds to the fitted empirical model described previously [Bost et al.]. **(B)** Effects of FoV size on the inferred *τ* parameter determined by empirical sampling (red), the negative binomial model (orange), and the beta binomial model (green). The dashed lines correspond to the fits to a power law. **(C)** Estimation of ⍺ using empirical sampling, the negative binomial model, and the beta binomial model. Vertical lines correspond to the estimated standard error when fit to the power law function. **(D)** Effects of **T** on the individual cell phenotype *–r* computed using the negative binomial model across various cell phenotypes. **(E)** Effect of FoV size on the probability of detecting 50 cells.

### Differential abundance testing is improved using negative and beta binomial distribution-based tests

Current common practice for comparison of abundances of a given cell phenotype between two groups of samples is to use a non-parametric rank test. Although the non-parametric approach limits false positive results, it has lower statistical power than adequate parametric tests and cannot provide effect size estimates or confidence intervals [Blair et al.]. We therefore compared the statistical power of the Wilcoxon rank test, the most commonly used rank-based test, to statistical tests based on negative and beta binomial regressions, namely the likelihood-ratio and Wald’s tests, respectively [Aban et al.].

In order to estimate the power of each test in a controlled fashion, we used the human lymph node dataset and performed a random thinning to simulate low cell density (Figure 3A). This approach resulted in point patterns for which the negative binomial model yielded a **µ** proportional to the fraction of thinned points, and the over-dispersion parameter **Θ** was unchanged compared to that for the original dataset (Figure S2A). The **p** and **ϕ** parameters of the beta binomial distribution model were proportional to the thinning rate (Figure S2B). As expected, based on point pattern theory [Illian et al. 2008], random thinning preserved spatial structures as shown by unaffected pair correlation functions (Figure S2C).

**Figure 3:**
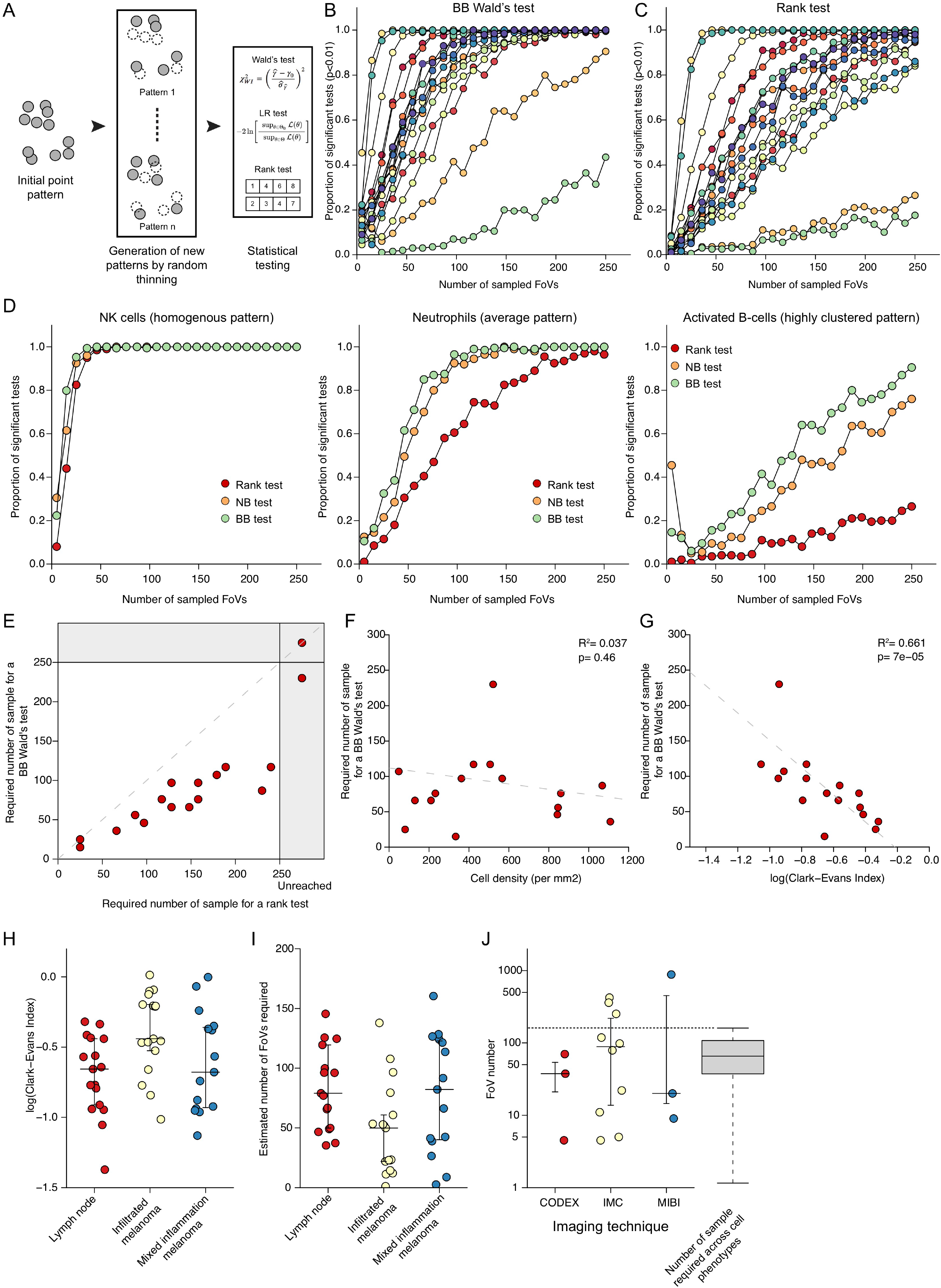
Differential abundance analysis of multiplexed imaging cell count data. **(A)** Illustration of workflow used to perform power analysis on cell count data. **(B & C)** Power analysis of B) the beta binomial-based Wald’s test and C) the Wilcoxon rank-based test. Each color corresponds to a specific cell phenotype. A log2 differential abundance of 1 and a p-value threshold of 0.01 were used. **(D)** Comparison of statistical powers of rank- and negative binomial- and beta binomial-based tests across three cell phenotypes with different spatial patterns. **(E)** Minimal number of FoVs necessary to detect significant changes in 80% of the simulations using a rank- or a beta binomial-based Wald’s test for each cell phenotype. Cell phenotypes for which the 80% threshold was not be reached with 250 FoVs were classified as ‘Unreached’. **(F)** Association between densities of individual cell phenotypes and required number of FoVs for a Wald’s test. The dashed line corresponds to a linear regression. **(G)** Association between Clark-Evans indices of individual cell phenotypes and required number of FoVs for a Wald’s test. The dashed line corresponds to a linear regression. **(H)** Clark-Evans indices for each individual cell phenotype across three samples. Horizontal bars correspond to the median and vertical bars to the IQR. **(I)** Estimated number of FoVs required for identification of each individual cell phenotype across three samples. Horizontal bars correspond to the median and vertical bars to the IQR. **(J)** Number of FoVs in published multiplexed imaging studies grouped based on technology: co-detection by indexing (CODEX), IMC, and multiplex ion beam imaging (MIBI). Horizonal bars correspond to the median and vertical bars to the IQR. To the right of the graph is the distribution of the estimated number of FoVs needed to detect a significant difference across cell phenotypes of all studied samples. The thick line corresponds to the median, and the bottom and upper limits of the box correspond to the first and third quartiles, respectively. The lower and upper whiskers correspond to the lowest and highest values, respectively, within the range of the first and third quartiles ±1.5 times the IQR.

We first measured how the sample size (i.e., the total number of FoVs) impacts the statistical power of each test to detect differential abundances of cell phenotypes. To do this, we generated two groups of samples, one through regular random sampling of FoVs, and another where half of the cells were randomly removed before performing random sampling, thus simulating a two-fold reduction of cell density for each cell phenotype. We observed considerable heterogeneity among cell phenotypes independently of the test used. Indeed, for some cell phenotypes, less than 20 samples resulted in a statistically significant test (>80% of test with a p-value lower than 0.01), whereas systematic statistical significance was not achieved even with hundreds of FoVs for other cell phenotypes (Figure 3B and C and Figure S2D).

We then compared the sensitivity of each statistical test across three representative cell phenotypes, NK cells, which have a homogenous point pattern (log2 Clark-Evans index = -0.33), activated B cells, which have a highly clustered point pattern (log2 Clark-Evans index = -0.94), and neutrophils, which have an intermediate pattern (log2 Clark-Evans index = -0.84). The three statistical tests gave similar results for the homogenous cell phenotype (Figure 3D, left panel). For the clustered phenotype, the beta binomial-based test was the most sensitive, followed by the negative binomial-based test, and finally the Wilcoxon rank test (Figure 3D, right panel). The same trend was observed for the intermediate phenotype, albeit with less pronounced differences (Figure 3D, center panel). A direct comparison of the minimal number of FoVs needed per samples to detect significant changes across all cell phenotypes present revealed that the beta binomial-based test required fewer FoVs than the Wilcoxon rank-based test or the beta binomial-based Wald’s test by up to two-fold (Figure 3E). We did not observe major differences when comparing Wald’s and likelihood-ratio tests for either beta or negative binomial distributions (Figure S2E and F).

We then sought to identify the factors that determine how many FoVs are needed to detect significant changes (>80% of test with a p-value lower than 0.01) for each cell phenotype. Density of the cells of the given phenotype did not significantly impact this value (Figure 3F, p=0.46), whereas spatial aggregation as measured by the Clark-Evans index was significantly correlated with the number of FoVs necessary (Figure 3G, R^2^=0.66, p=7.0e-5).

This correlation prompted us to compute the Clark-Evans index across cell phenotypes in our lymph node and two melanoma samples (Figure 3H). We then inferred the associated number of FoVs necessary to detect significant differences by training a model on the lymph node simulations. We observed that in the two melanoma tissue samples, between 1 and 160 FoVs are required to confidently detect statistical differences, depending on the cell phenotype (Figure 3H and I). For highly aggregated cell types most previously published multiplexed imaging studies would have been underpowered without pre-selecting FoVs as they have less than 150 FoVs per group (Figure 3I and J). In sum, binomial distribution-based statistical tests outperform the commonly used Wilcoxon rank test for comparison of cell densities of non-homogenously distributed cell types between two groups of samples. In addition, statistical power strongly varies across cell phenotypes and is driven by cell spatial pattern and not cell density.

### Stratified sampling drastically decreases cell density estimation variance

Due to sample availability (e.g. patient needle biopsy), experimental complexity and cost, analysis of high sample numbers is not always attainable, imposing strong limitations on the statistical analysis, especially for aggregated cell phenotypes. We therefore investigated whether more complex sampling strategies could overcome this challenge. We implemented a stratified sampling approach [Cochran] that we call Spatial Stratified Sampling. Stratified sampling works in a two-step manner: first, members of a sample population are split into groups called strata that are assumed to be homogeneous before performing random sampling of each stratum. Individual estimates across strata are then merged to provide a final estimate for the whole population. Usually, samples are split using an easy-to-measure variable, such as geographic location for political polls. When groups are homogenous, stratified sampling dramatically reduces the variance of the estimator (i.e. the function that is used to estimate an unknown parameter based on measured data) compared to random sampling, allowing measurement of fewer samples without loss of quality. In the case of Spatial Stratified Sampling, a pan-tissue image such as an immunofluorescence image obtained by a slide-scanner, of a marker (either protein or transcript) is used to stratify the tissue into distinct regions. Then, random sampling is performed within each individual stratum to estimate the cell density of each cell phenotype across the strata before computing the final cell density estimation (Figure 4A).

**Figure 4:**
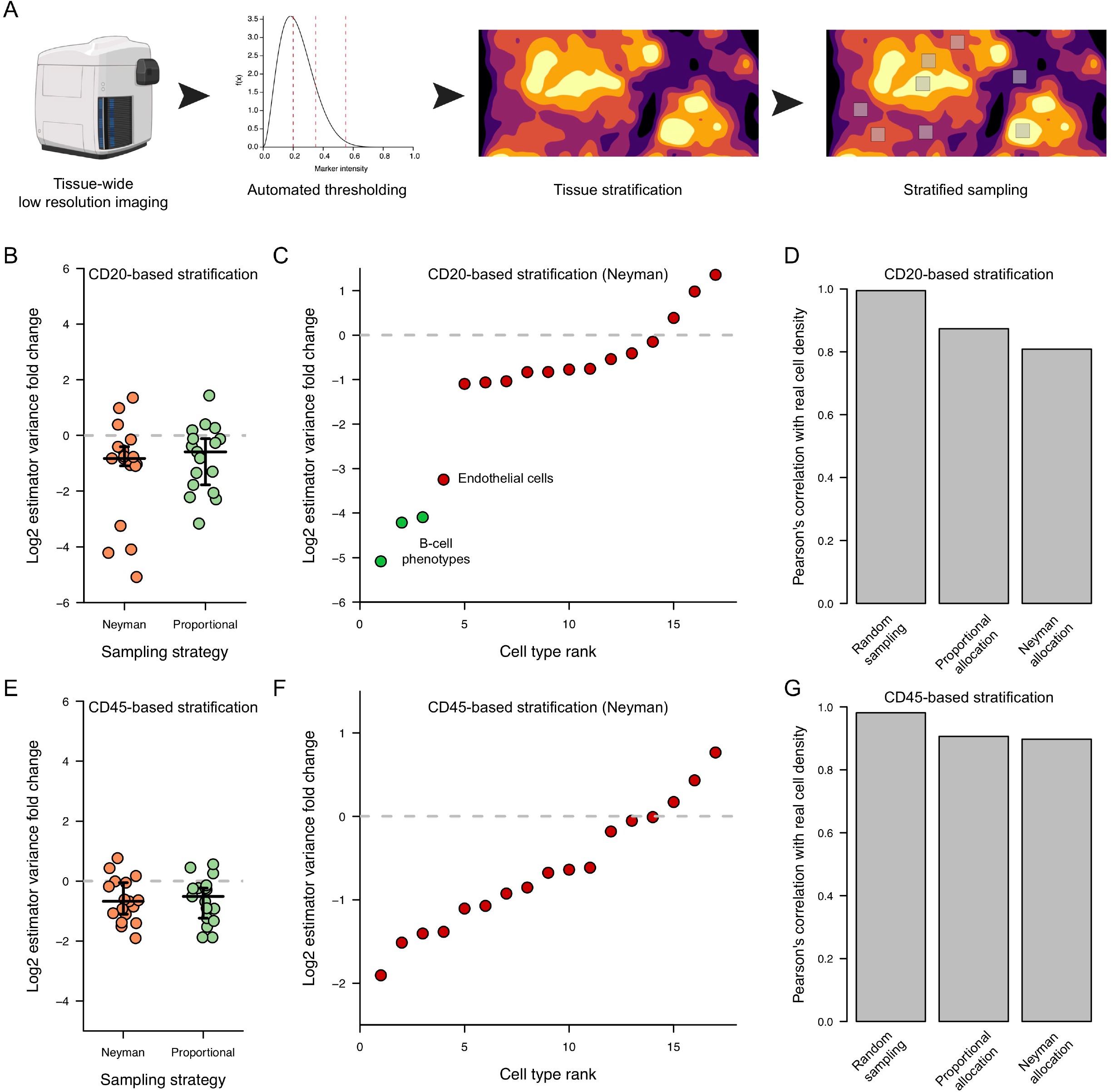
Use of stratified sampling improves robustness of cell density estimation. **(A)** Stratified sampling workflow for multiplexed imaging. **(B)** Log2 ratio between the cell density variance estimator obtained by stratified and random sampling for the lymph node sample data. CD20 was used to perform the stratified sampling. Each dot corresponds to one cell phenotype. **(C)** Ranked log2 ratio between the cell density variance estimator obtained by Neyman stratified and random sampling for the lymph node sample data. CD20 was used to perform the stratified sampling. **(D)** Correlation between estimated and sampled cell densities obtained using CD20 for the stratified sampling of the lymph node sample data. **(E)** Log2 ratio between the cell density variance estimator obtained by stratified and random sampling for the lymph node sample data. CD45 was used to perform the stratified sampling. Each dot corresponds to one cell phenotype. **(F)** Ranked log2 ratio between the cell density variance estimator obtained by Neyman stratified and random sampling for the lymph node sample data. CD45 was used to perform the stratified sampling. **(G)** Correlation between estimated and sampled cell densities obtained using CD45 for the stratified sampling of the lymph node sample data.

To test our approach, we used our lymph node dataset and a generated a simulated CD20 immunofluorescence image by applying a Gaussian blur to the original image. We selected the CD20 marker as it delineates lymph node anatomical zones and might therefore improve the estimation of the highly clustered B cell phenotype density. We stratified the imaged tissue into six strata based on CD20 intensity using Minimal Variance Stratification thresholding and then simulated two types of stratified samplings, proportional and Neyman, with 10 square FoVs of 100 µm width.

The variance of the cellular density estimator was significantly lower for both stratified sampling methods than for random sampling (Figure 4B, univariate t-test, p=0.010 and p=0.011 for proportional sampling and Neyman sampling, respectively). Moreover, for the three B cell phenotypes, estimator variance was considerably lower when Neyman sampling was used (by at least 16-fold) than when random sampling was used (Figure 4C). This indicated that CD20-based stratification was very efficient on CD20-expressing cells. This variance reduction came at the cost of an increased bias, however, which is a common issue in supervised learning. This is illustrated by the decreased correlation of the density estimate for all types of sampling compared with the known density (Figure 4D). We then investigated whether changing the stratifying marker impacted the quality of Spatial Stratified Sampling. When the pan-immune cell marker CD45 was used as the covariate, estimator variance was reduced for most cell phenotypes compared to random sampling (Figure 4E univariate t-test, p=2.0e-3 and p=8.0e-4 for proportional and Neyman sampling, respectively). No cell phenotype had a variance reduction factor larger than 4, even with Neyman stratification (Figure 4C and D). Use of CD45 also resulted in a lower bias than did use of CD20 (Figure 5G), suggesting that the stratified sampling is indeed subject to bias-variance tradeoff.

To further evaluate the robustness of the Spatial Stratified Sampling approach, we applied the strategy to melanoma samples using either the pan-immune marker CD45 or HLA-DR, which is expressed only on antigen presenting cells, as stratification markers. For the melanoma sample with spatially homogenous immune cell infiltration, we observed that stratification with either marker resulted in less variable cell density estimators than random sampling (Figure S3A and B). Interestingly, only the CD45-based stratification reduced the estimator variance by a factor higher than 10, and this only for two B cell phenotypes (CD20-positive cells). We observed a similar pattern when stratified sampling was performed on the second melanoma sample, which has a heterogeneous immune infiltration pattern (Figure S3C and D). For this sample, with higher variance reduction was achieved using CD45 than HLA-DR, suggesting that CD45 is a better stratification marker than HLA-DR for melanoma samples independently of their immune infiltrate status. In summary, stratified sampling improves the precision and robustness of multiplexed imaging analysis and should be applied specifically to highly aggregated cell phenotypes that cannot be studied using regular random sampling.

## Conclusion

Here we reported the first comprehensive description of the statistical properties of multiplexed imaging cell count data within a single FoV using simple, yet efficient, probabilistic negative binomial distribution and the beta binomial distribution models. Moreover, we determined the precise effects of FoV size on both the mean and over-dispersion parameters of both distributed and linked cell phenotype spatial patterns to over-dispersion parameters. From these two models, we built new differential abundance statistical tests that provided a considerable gain of statistical power compared to commonly used non-parametric rank-based tests. In addition, our analysis revealed that the statistical power of a multiplexed imaging experiment is dependent on the spatial segregation of a given cell phenotype. Some cell count data display a high variability due to the high spatial segregation of their associated cell phenotypes, and random sampling cannot provide a robust measurement of cell density, making differential abundance analysis extremely challenging. To overcome this challenge, we introduced the use of stratified sampling for analysis of multiplexed imaging data. Stratified sampling drastically enhanced the robustness of cell density estimates, especially for highly clustered cell phenotypes, and improved the statistical power of multiplexed imaging-based studies. Use of the Spatial Stratified Sampling approach will allow these technologies to be used to analyze highly complex diseases and when few samples or only few FoVs are available/can be measured.

It should be noted that our modeling approach was not designed for the analysis of spatial transcriptomic datasets such as those generated by the Visium® platform [Janesick et al.]. As spatial transcriptomics experiments become more widely used, appropriate computational methods to analyze the generated data will become crucial. However, spatial transcriptomic data strongly differ from multiplexed imaging data, as in spatial transcriptomics datasets individual points are localized on a predefined dense lattice and do not correspond to individual cells due to the current spatial resolution (∼50 µm). Therefore, future work should address these limitations and extend the range of application of our model.

Stratified sampling of multiplexed imaging data can drastically improve the statistical power of differential density analysis compared to random sampling. However, additional work will be required to make it applicable for use on a large scale. First, the potential statistical power gain for more complex analysis, such as survival analysis or cell-cell interaction analyses, common analyses when studying cancer tissue sections [Jackson et al., Schürch et al.], need to be investigated. Secondly, an alternative to the cumbersome requirement of jointly performing a whole tissue-section immunofluorescence and IMC experiment should be found. An alternative approach would be to use IMC or MIBI [Moffitt et al.] to image a limited set of discontinuous pixels over a large area to then to computationally reconstruct a low-resolution image [Achanta et al].

Spatial analysis of clinical samples is usually performed by immunohistochemical or H&E staining. Those methods only measure a very limited set of features but can be performed at low cost and provide information on the whole tissue section, allowing their clinical use in a routine manner. Multiplexed imaging approaches follow a reverse strategy: these technologies image tens of features on a limited area of the tissue section. Multiplexed imaging techniques are currently not used clinically, but the development of appropriate and improved statistical methods, such as those presented in this article, could pave the way toward routine use of such powerful imaging techniques. The methods we present are time and cost efficient and will significantly improve multiplexed imaging experiment efficiency.

## Methods

### Melanoma section processing and IMC data acquisition

Melanoma FFPE blocks were obtained from the University Hospital Zurich (KEK-ZH-Nr 2014-0425), and 4-µm thick sections were collected onto Adhesion Superfrost® Plus slides and placed in an oven for 30 min at 70 °C before being stored at -20 °C until staining. Sections were dewaxed, rehydrated, and subjected to a heat-induced epitope retrieval step for 30 min at 95 ºC in 10 mM Tris, pH 9.2, 1 mM EDTA. The sections were then incubated in blocking buffer (3% BSA in TBS-T) for 1 h at room temperature before incubation with a 27-antibody panel (Supplementary Table S1) diluted in blocking buffer overnight at 4 ºC. Nuclear staining was then performed by adding an iridium solution (5 nM) diluted in TBS (1:100 dilution) to the sample and incubating for 5 min. The samples were then washed three times (10 minutes per wash) in TBS and dried. Images were acquired using a Hyperion Imaging System with the ablation frequency set to 200 Hz and the ablation energy set to 6 dB with X and Y steps set to 1 µm.

### IMC data pre-processing and analysis

The raw mcd files were processed using the Steinbock pipeline, version 0.13.4 [Windhager et al.]. In brief, the raw files were converted into tiff files, and the cells were segmented using a pre-trained neural network [Greenwald et al.] using H3K9ac as the nuclear channel and Vimentin and CD45 as the membrane channels. Default parameters were used for Mesmer [Greenwald et al.]. The mean channel intensity was then computed for each cell and exported as a text file, together with the location, the size, and other basic information on the cells. The single-cell IMC data were then analyzed using in-house R scripts (R version 4.0.3). Each channel was normalized by performing a Poisson regression between the total channel intensity and the cell size (in pixels); the Pearson’s residuals were extracted as the new scaled values. The cells were then clustered by first building a *k*-nearest-neighbor graph with 15 neighbors (using cosine distance) and then clustered using Louvain’s community detection implemented in the **igraph** package with default parameters.

### Binomial distribution fitting and analysis

The negative binomial distribution was fitted using the fitdist() function from the **fitdistrplus** R package using the moments method. The beta binomial distribution was fitted using the betabin() function from the **aod** R package where both the probability of success and the over-dispersion parameter were considered to be constant. Linear modeling between parameters and FoV width was done using the lm() R function.

### Computation of the AIC score

The AIC score of a given model is defined as:

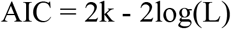

where k is the number of parameters of the model, and L the model likelihood. k was 1 for the Poisson model and 2 for the binomial models. Log likelihood was computed by assuming that each observation is independent and summing individual observation log-likelihoods.

### Estimation of *α* and *τ* parameters

Theoretical curves describing the expected number of cell phenotypes to be recovered when imaging a given number of FoVs were obtained by computing the probability mass function of the sum of ***n*** random variables of fitted parameters for each cell phenotype and then computing the probability of observing at least **T** cells. Here **T** was set to 50 as described previously [Bost et al.]. In the case of the negative binomial distribution, we used the fact that the distribution is stable, allowing us to directly compute the probability mass function of the new random variable that is also a negative binomial variable (Appendix 1). For beta binomial variables, the probability mass function was computed through iterative convolution using the convolve() R function. Once the probability of recovering each cell phenotype individually was computed for a given number of FoVs, the expected total number of recovered cell types was obtained by summing values across all cell phenotypes.

### Random thinning of point patterns and associated analysis

For each cell phenotype, we randomly removed a given proportion of cells using the sample() function from the base package without replacement and then performed random FoV sampling using a width of 400 µm. Binomial distributions were then fitted as described above. This was repeated 50 times for different proportions of removed cells (10% to 90% cells removed). The relationship between the proportion of thinned cells and the mean estimated parameter values was then calculated using the lm() function (**stats** package) associated p-values were calculated using the anova() function from the **stats** package. The pair correlation function was estimated using the Kest() and pcf() functions from the **spatstat** R package. Kest() was used using default parameters, whereas the pcf() function was used with the parameter ‘method’ set to ‘b’. An envelope was plotted with the upper and lower limits of the envelope representing the standard deviation.

### Differential abundance analysis

Differential abundance analysis based on a negative binomial model was performed using the glm.nb() function from the **MASS** R package where the group to which the sample belongs was used to predict the individual **µ** values. A logarithmic link function was used. The beta binomial-based analysis was performed using the betabin() function from the **aod** R package. Individual **p** parameters were predicted using the sample group as a covariate and the logistic function as link function. Individual **ϕ** values were also computed using the sample group as a covariate with identity link function.

The p-values associated with the negative binomial-based likelihood ratio test were computed using the anova.negbin() function from the **MASS** package. For the negative binomial-based Wald’s test, the estimated *γ* parameter and its variance were extracted from the fitted model, raised to the power of 2, and the ratio was used to perform a one-sided chi^2^ test. The p-values associated with the beta binomial-based likelihood ratio test were computed in a two-step manner. First, two models were fitted, one using the sample group as a covariate and the other without. Then the likelihood ratio tests were performed using the anova() function from the **aod** package. The p-values for the beta binomial-based Wald’s test were computed using the wald.test() and vcov() functions from the **aod** package. Rank-based tests were performed using the kruskal.test() R function.

### Inference of the minimum number of samples for statistically powerful experiments

To infer the number of samples required to detect a significantly different abundance between two samples for a given cell phenotype, we used the strong relationships between the number of FoVs required and Clark-Evans index. To predict the minimum required number as a function of the Clark-Evans index, we did not use a simple linear regression as a minimal number cannot be negative. Instead, we used the following model:

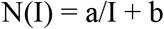

Where **N()** is the minimal number of FoVs, **I** is the Clark-Evans index, and **a** and **b** are two positive variables. This model was fitted using the nls() function with default parameters with 1 as starting values for **a** and **b** parameters.

### Simulation of immunofluorescence images and stratification

Smoothing of the raw IMC images was done using Python 3.8 and the **sci-kit image** package. Briefly, raw IMC tiff files were loaded and pixel intensity was rescaled to the (0,1) range and then blurred using the gaussian() function with a sigma parameter set to 50. Values were then min-max scaled to be contained between 0 and 1. Resulting images were then saved in the tiff format. Stratification of the image was done using the Dalenius approximation of the minimal variance stratification approach [Dalenius et al.]. We assumed that the distribution of the smoothed image pixel intensities followed a beta distribution (due to the scaling) and computed the corresponding minimal variance stratification threshold. We let f be the probability density function of the variable to sample, or of a highly correlated variable. Here the blurred intensity of the marker of interest is considered as the variable of interest. We defined y(u) as:

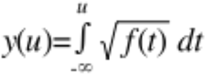

The minimal variance stratification approach consists of identifying the L thresholding values {x_1_, x_2_,…, x_L_} by solving the equations:

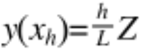

with

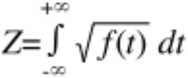

In the case of a beta distribution with parameters ⍺ and β:

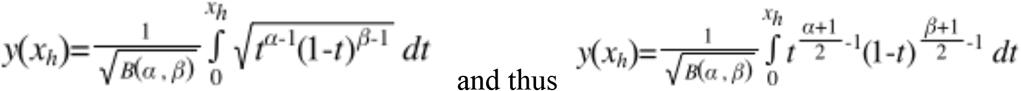

where B(,) is the complete beta function. Therefore:

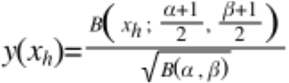

where B(;,) is the incomplete beta function. The normalization constant Z is described by the following equation:

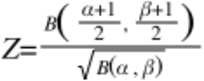

And, thus:

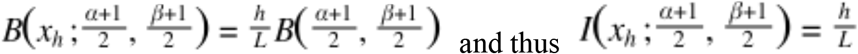

where I(;,) is the regularized incomplete beta function. The last equation was numerically solved through least square minimization by the optimize() R function with default parameters. The regularized incomplete beta function values were computed using the Rbeta() function from the **ZipfR** package. Parameters of the beta distribution were inferred using the fitdist() function of the **MASS** package by the moment method. The tiff files were loaded into R using the readTIFF() function from the **tiff** package.

### Stratified sampling

In order to sample ***n*** FoVs of width *w* from a given stratum, we used the following approach. We first computed the L0 distance transform map of the binarized image (i.e., the pixel part of the stratum) using the distance_transform() function from the **imager** package. We then randomly selected a pixel with a value larger than *w*/2 using the normalized pixel value as sampling probability. Next, a new L0 distance transform map was computed using the original binarized image with the values of pixels within the sampled FoV set to 0. A new FoV was then sampled, and the binary image was updated until all ***n*** FoVs were sampled or until no new FoV could be sampled (i.e., no pixels within the stratum were further than *w*/2 from another strata).

The number of FoVs sampled in each stratum for the proportional stratified sampling was computed by first dividing the surface area of each stratum by the total sample area and then multiply the obtained value by the total number of FoV before rounding the final value. The obtained vector was changed if one stratum had no FoVs; in this case, one FoV was removed from the largest stratum and was assigned to stratum without an FoV.

### Estimation of the cell count variance for Neyman allocation

The Neyman allocation requires an estimation of the variance of cell density across strata. Although this is not feasible in practice, we made this estimate based on Poisson count model. We assumed that the intensity of the marker of interest in a FoV is proportional to the number of cells of interest within this FoV. As the number of cells per FoV is over-dispersed, the variance of the number cells is at least equal to the mean number of cells. Therefore, the variance of cell count can be considered as being proportional to the mean marker intensity. The number of FoV per stratum was then computed similarly to the proportional sampling except that the stratum area proportion was multiplied by the estimated standard deviation before multiplying by the total number of FoV and rounding.

## Code availability

All scripts and functions developed for this paper are available in the **Balagan** R package available on GitHub (https://github.com/PierreBSC/Balagan).

## Acknowledgements

We acknowledge Bodenmiller lab members for critical reading and providing feedback on the manuscript and thank Alice Balfourier and Ronan Thibaut for their valuable advices. P.B. is funded by an EMBO postdoctoral fellowship (fellowship number ALT 427-2021).

## Author Contributions Statement

P.B. developed the methodology, implemented the R code, performed the IMC experiments and wrote the manuscript. R.C. and U.M. developed the methodology and wrote the manuscript. B.B. oversaw the project, assisted with study design, acquired funding, and helped write the manuscript.

## Competing Interests Statement

The authors declare no competing interests.

## Figure Legends

**Supplementary Figure 1:**
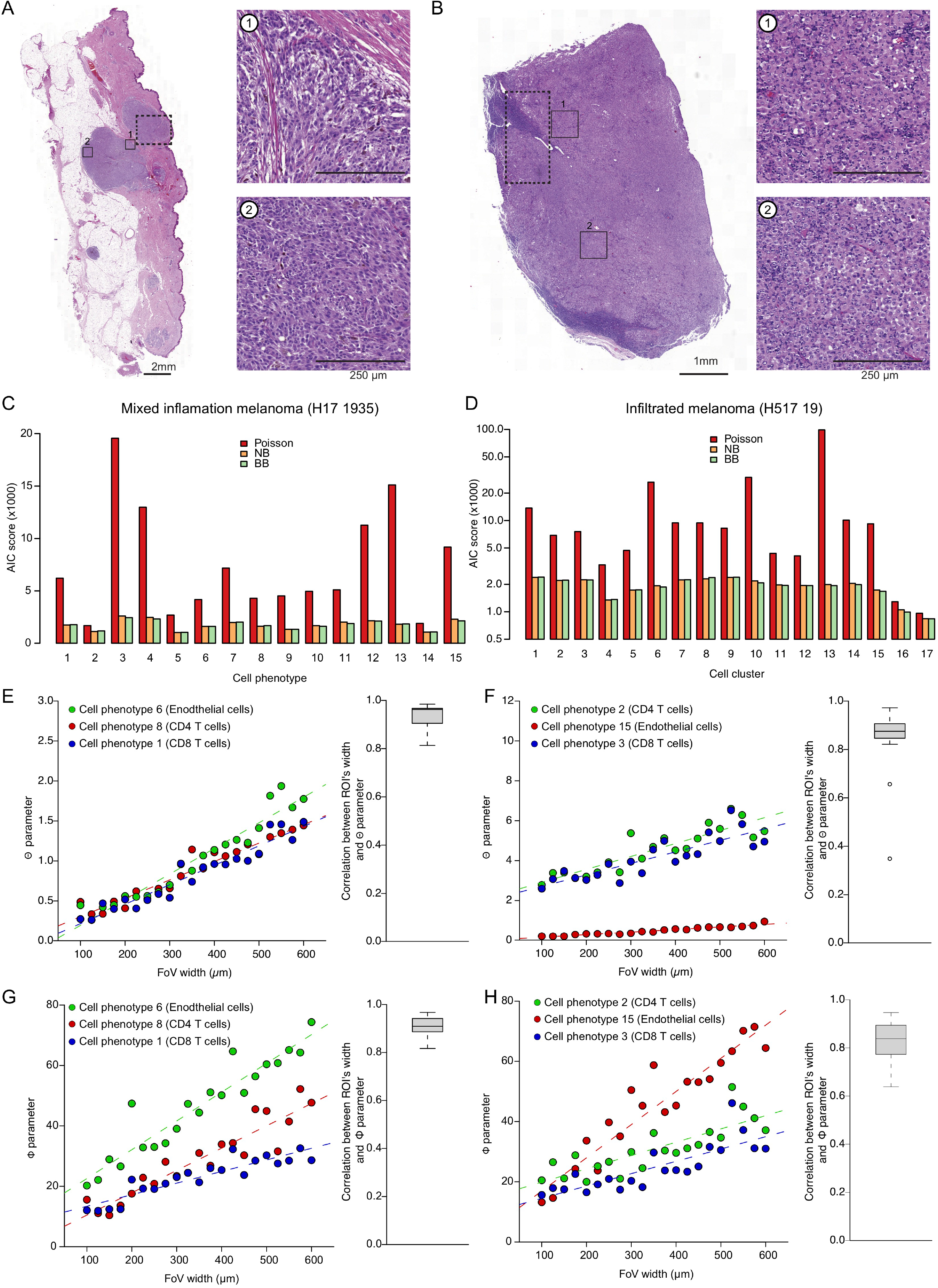
**(A & B)** H&E stained sections of melanoma samples with A) spatially inhomogeneous and B) spatially homogeneous immune cell infiltration. Small rectangles with plain borders indicate regions shown at higher magnification to the right. Large rectangles with dashed borders correspond to regions imaged by IMC. **(C & D)** AIC scores for each model for each cell phenotype for C) inhomogeneous sample and D) the homogeneous sample. **(E & F)** Left: Relationship between FoV width and negative binomial **Θ** for three cell phenotypes of the melanoma sample with E) inhomogeneous immune cell infiltration and F) homogeneous immune cell infiltration. Dashed lines correspond to the linear model between FoV width and **Θ**. Right: Distribution of R^2^ values for the linear relationship between FoV width and **Θ** parameter across the cell phenotypes. **(G & H)** Left: Relationship between FoV width and beta binomial parameter **ϕ** for three cell phenotypes of the melanoma samples with G) inhomogeneous and H) homogeneous immune cell infiltration. Dashed lines correspond to the linear model between FoV width and **ϕ**. Right: Distribution of R^2^ values for the linear relationship between FoV width and **ϕ** across the cell phenotypes.

**Supplementary Figure 2:**
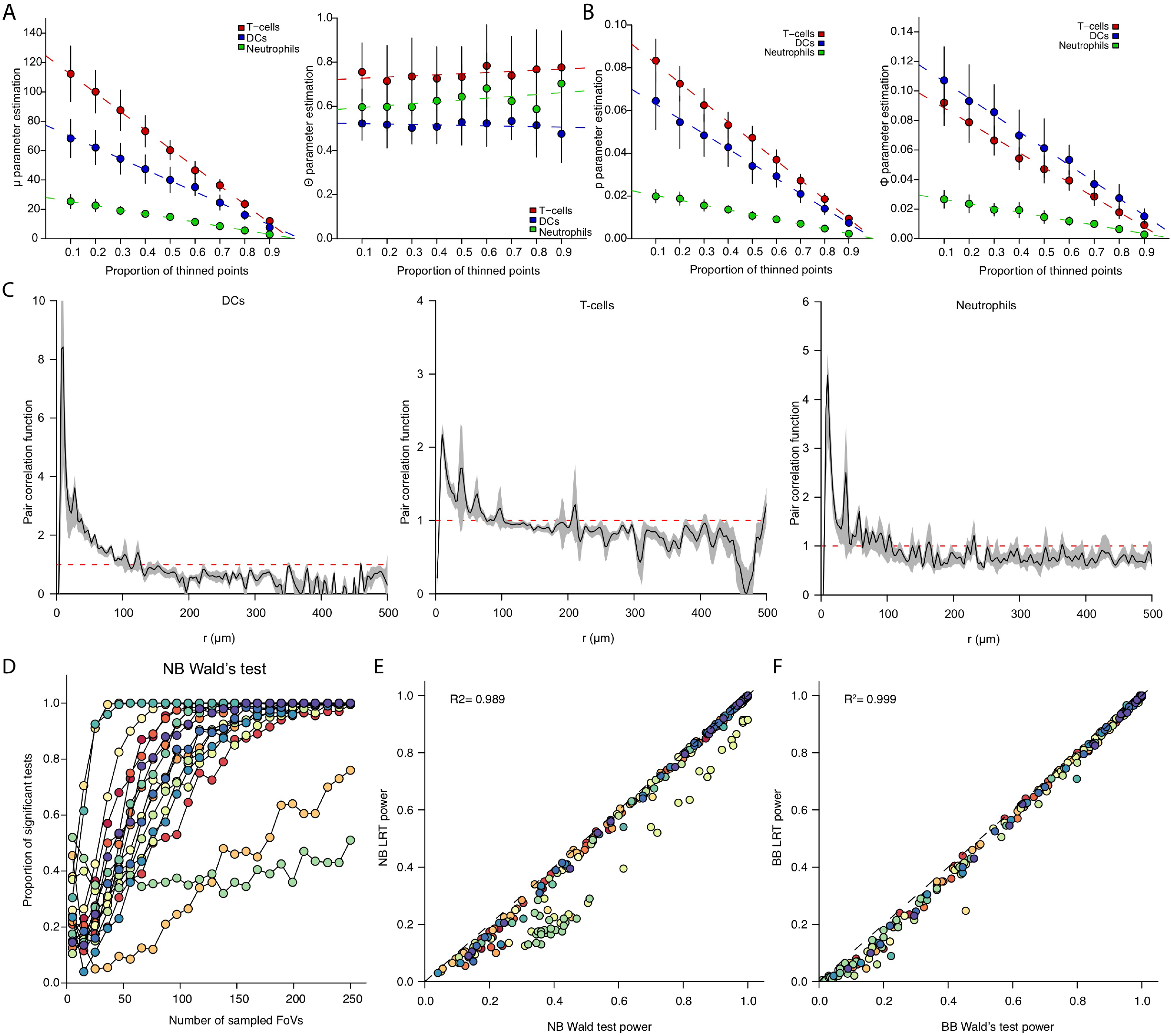
**(A)** Effects of random point thinning on estimated **µ** (left) and **Θ** (right). Three representative cell phenotypes are shown. Dashed lines correspond to the linear regression between the proportion of thinned points and estimated parameter values. Vertical lines correspond to the standard deviation observed across samplings. **(B)** Effects of random point thinning on estimated **p** (left) and **ϕ** (right). Dashed lines correspond to the linear regression between the proportion of thinned points and estimated parameter values. Vertical lines correspond to the standard deviation observed across samplings. **(C)** Effects of the random thinning on the point pattern structure. The pair correlation function was computed for three cell phenotypes before (black line) or after (grey envelope) random thinning. The envelope’s border corresponds to the mean ± the standard deviation observed across the simulations. **(D)** Power analysis of the negative binomial-based Wald’s test. Each color corresponds to a specific cell phenotype. A log2 differential abundance of 1 and p-value threshold of 0.01 were used. **(E)** Comparison of the statistical power of negative binomial-based Wald’s and likelihood ratio tests. A log2 differential abundance of 1 and p-value threshold of 0.01 were used. Each dot corresponds to a defined number of FoVs and cell phenotype. Dots are colored according to their associated cell phenotype. **(F)** Comparison of the statistical power of beta binomial-based Wald’s and likelihood ratio tests. A log2 differential abundance of 1 and p-value threshold of 0.01 were used. Each dot corresponds to a defined number of FoVs and cell phenotype. Dots are colored according to their associated cell phenotype.

**Supplementary Figure 3:**
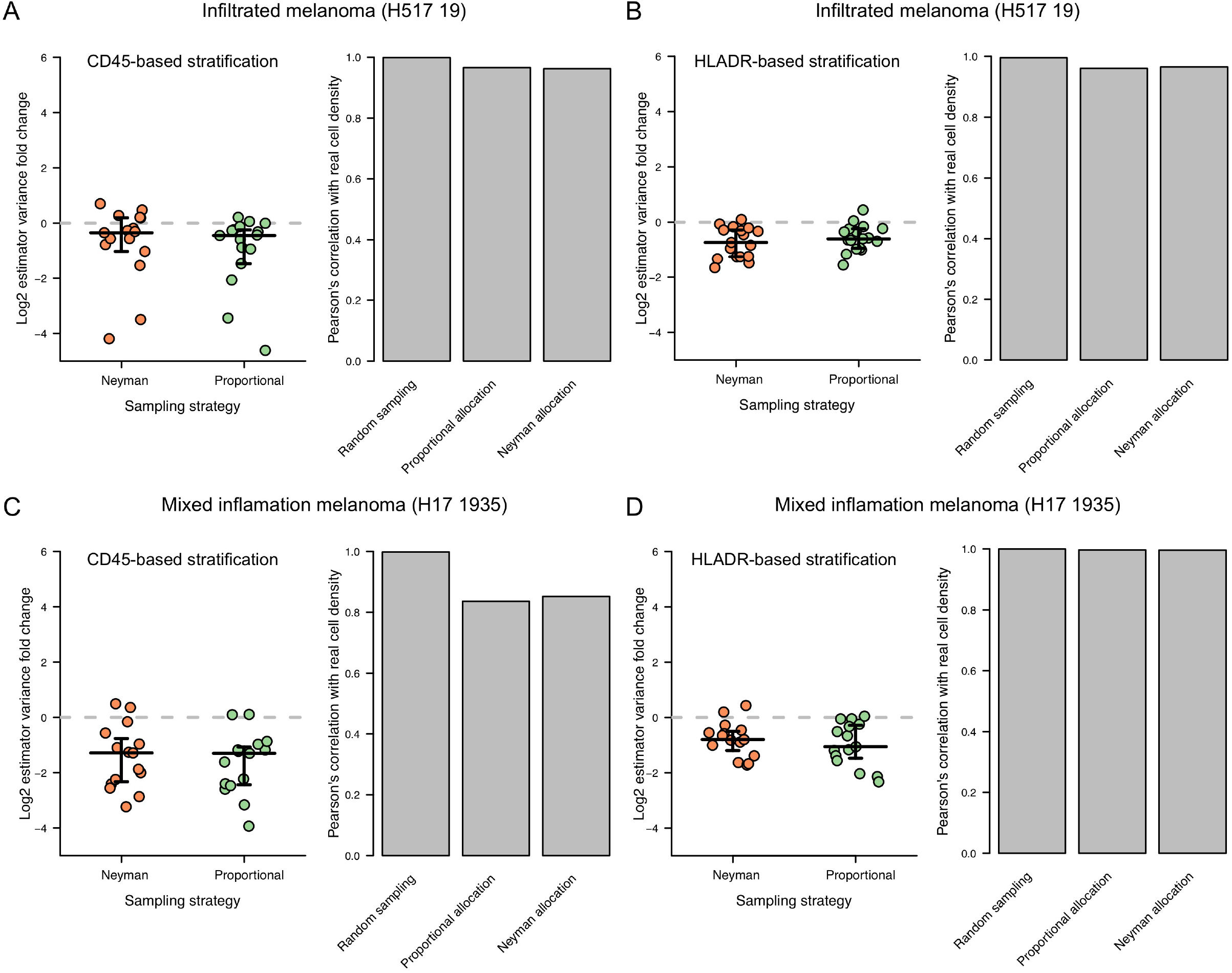
**(A & B)** Left: Log2 ratio between the cell density variance estimator obtained by stratified sampling with A) CD45 and B) HLA-DR and random sampling for the melanoma sample with homogeneous immune cell infiltration. Each dot corresponds to one cell phenotype. Right: Correlation between estimated and sampled cell densities. **(C & D)** Left: Log2 ratio between the cell density variance estimator obtained by stratified sampling with C) CD45 and D) HLA-DR and random sampling for the melanoma sample with inhomogeneous immune cell infiltration. Each dot corresponds to one cell phenotype. Right: Correlation between estimated and sampled cell densities.

## Notes

### Competing Interest Statement

The authors have declared no competing interest.

